# White matter haemodynamics: basic physiology and disruption in neuroinflammatory disease

**DOI:** 10.1101/208751

**Authors:** Scott. C. Kolbe, Sanuji. I. Gajamange, Jon. O.S.H. Cleary, Trevor. J. Kilpatrick

## Abstract

The white matter is highly vascularised by the cerebral venous system. The neuroinflammatory disease multiple sclerosis is associated with infiltration of peripheral immune cells into the brain via these vessels. Understanding venous pathophysiology in multiple sclerosis is thus critical for understanding early disease aetiology. In this paper, we describe a unique blood oxygen-level dependent (BOLD) signal within the white matter using functional MRI and spatial independent components analysis, a blind signal source separation method. The signal was characterised by a narrow peak frequency band between 0.05 and 0.1 Hz. Hypercapnia (transient breath holds), known to alter venous calibre in cortex, induced transient increases in white matter BOLD that disrupted the oscillation indicative of a vasodilatory/contractile mechanism. Comparison of the white matter BOLD oscillations between age and sex matched groups of 18 multiple sclerosis and 14 healthy participants revealed a loss of power in the white matter BOLD signal in the peak frequency band (patients = 6.70±0.94 dB/Hz vs controls = 7.64±0.71 dB/Hz; p=0.006). In multiple sclerosis patients, lower power was associated with greater levels of neuroinflammatory activity (R=−0.64, p=0.006) but not neurodegenerative disease markers. Using a signal modelling technique, we assessed the anatomical distribution of white matter BOLD signal abnormalities and detected reduced power in the periventricular white matter, a region of known venous damage in multiple sclerosis patients. These results demonstrate a novel link between neuroinflammation and vascular physiological dysfunction in the cerebral white matter, and could indicate enduring loss of vascular compliance associated with imperfect repair of blood-brain barrier damage after resolution of acute neuroinflammation.

## Introduction

Multiple sclerosis is a chronic inflammatory disorder of the CNS. The initiating neuroinflammatory events of multiple sclerosis remain unclear, however, the pathological hallmark of the disease involves peripheral immune cell infiltration via the internal cerebral venous system of the white matter (Tan et al., 2000) upon disruption to the blood-brain-barrier, leading to perivenular demyelination and axonal injury.

Despite the relevance of the white matter venous system to multiple sclerosis pathology, relatively few studies have directly studied venous pathology in the multiple sclerosis brain. Pathological studies have reported enduring damage to the venous system in multiple sclerosis after the resolution of acute inflammation (Adams, 1988; Adams et al., 1985). Neuroimaging studies have reported reduced density (Ge et al., 2009; Sinnecker et al., 2013; Zivadinov et al., 2011), calibre (Gaitan et al., 2013), and perfusion (Varga et al., 2009) of the white matter venous system in multiple sclerosis patients. Together, these findings indicate that the influx of peripheral inflammatory cells into the brain via the cerebral venous system leads to enduring anatomical and physiological changes to the vessels. It is conceivable that such pathologies could confer a susceptibility to further acute inflammation or lead to hypo-perfusion, thus exacerbating neuronal injury.

To further investigate physiological changes in the cerebral venous system, we assessed the blood oxygen-level dependent (BOLD) signal in the white matter using functional MRI. We show that the cerebral venous system is characterised by a unique oscillatory BOLD signal that is disrupted in people with early multiple sclerosis, and that the degree of signal disruption is associated with the degree of neuroinflammation.

## Materials and Methods

### Subjects

Eighteen participants (7m/9f; mean (SD) age at the time of testing = 42.4 (10.3) yrs) were recruited prospectively between 2008 and 2010 upon presenting with first demyelinating event acute optic neuritis at the Royal Victorian Eye and Ear Hospital as part of a published trial investigating optic nerve imaging during and after acute optic neuritis (van der Walt et al., 2013). Scanning for the present study was performed between 3 and 5 years after initial recruitment as part of an extension study. At the time of scanning, all patients had been diagnosed with clinically definite multiple sclerosis based on the 2010 revisions to the McDonald criteria (Polman et al., 2011). Fourteen control subjects (6m/9f; mean (SD) age at the time of testing = 32.5 (4.6) yrs) were also recruited. This study was approved by the Royal Victorian Eye and Ear Hospital and Royal Melbourne Hospital Human Research Ethics Committees. All participants provided voluntary written consent in accordance with the Declaration of Helsinki.

### MRI acquisitions

All MRI scans were performed using a 3 Tesla MRI system (Trio TIM, Siemens, Erlangen, Germany) with a 32-channel receiver head coil. Three MRI sequences were acquired for each subject: (1) A 3D whole brain double inversion recovery sequence for lesion identification (repetition/echo/inversion times = 7400/324/3000 ms; flip angle = 120°; resolution 0.55×0.55×1.1 mm^3^ sagittal acquisition); (2) A 3D whole brain T1-weighted sequence for volumetric assessments (repetition/echo/inversion times = 1900/2.63/900 ms; flip angle = 9°; resolution 0.8×0.8×0.8 mm^3^ sagittal acquisition); and (3) A BOLD-weighted echo planar imaging sequence acquired while subjects watched a blank screen with eyes open (repetition/echo times = 1400/33 ms; flip angle = 85°; resolution 2.5×2.5×2.5 mm^3^ axial acquisition; number of BOLD measurements = 440). High spatial and temporal resolution was obtained through the use of “multi-band” simultaneous multi-slice echo planar imaging acquisition (3× acceleration) (Xu et al., 2013) and GRAPPA in-plane acceleration (2× acceleration). In addition, a single-band image was acquired for each subject (no multi-band acceleration) which, because of its enhanced grey/white matter contrast, was used for registrations between echo planar imaging and T1 anatomical space.

Lesions were delineated on the double inversion recovery images using a semi-automated thresholding technique (Rorden and Brett, 2000). Intra-cranial, whole brain parenchyma, grey matter and white matter volumes were obtained using Freesurfer (version 5.0.4) and the standard analysis pipeline (Dale et al., 1999). These measures were used to calculate brain, grey matter and white matter volumes, normalised to the intra-cranial volume for each subject.

### BOLD MRI pre-processing and spatial independent components analysis

BOLD-weighted MRI scans were processed using FSL MELODIC spatial-ICA software (Beckmann and Smith, 2004). The processing pipeline was as follows for each subject. (1) BOLD time-series image data were linearly realigned and registered to the single-band image. One patient and one control subject were excluded due to severe head motion (>1 mm inter-scan motion) during BOLD imaging. (2) Data were high pass filtered (pass band > 0.01 Hz) to remove low frequency signal drift due to MRI gradient heating and spatially smoothed using a non-linear edge-detection based algorithm (Smith and Brady, 1997) with spatial extent of 5 mm. (3) Spatial ICA was performed on each subject.

The resulting independent component maps were manually inspected and the white matter component was identified for each subject based on two anatomical features: (a) within the white matter, and (b) in close proximity to the lateral ventricles. Potential differences between patients and controls in the spectral qualities of the independent component time-courses (peak power and frequency) were tested for using Student’s t-tests. Pearson’s correlation analyses were used to test for co-variation between peak power and frequency.

Independent component maps were transformed to standard MNI-152 brain space using a three stage non-linear registration procedure utilising the Advanced Normalisation Tools (ANTs) software (Avants et al., 2014). (1) Each subject’s echo planar imaging reference image was nonlinearly registered to the subject’s T1 anatomical image. (2) Each subject’s T1 anatomical image was registered to the MNI-152 T1 standard brain. (3) The two deformation fields were added together and the independent component maps were transformed to MNI space using the resulting field. Example venous independent component maps in standard MNI space are shown for two patients and two control subjects in Figure 1A. Potential differences in the anatomical distribution of the IC maps between patients and controls was tested using voxel-wise unpaired t-tests with significance level calculated FSL’s RANDOMISE non-parametric permutation sampling statistical tool (Winkler et al., 2014) corrected for multiple comparisons using the threshold-free cluster enhancement algorithm (Smith and Nichols, 2009).

To generate a probabilistic standard space map of the white matter veins based on the independent component maps, each subject’s standard space independent component map was thresholded (Z > 2.3) and binarised. The resulting binary masks were averaged across all subjects to obtain a map where each voxel value represented the proportion of subjects where that voxel was within the venous independent component map (Figure 3A).

### Replication using Human Connectome Project Data

Thirty random healthy resting-state 3 Tesla fMRI datasets were obtained from the Washington University – University of Minnesota Human Connectome Project (Van Essen et al., 2013). Prior to downloading, the datasets had been minimally pre-processed prior to downloading according to the protocol described in (Glasser et al., 2013) including motion correction and nonlinear registration to MNI-152 atlas space. After downloading we performed spatially smoothed using a non-linear edge-detection based algorithm (Smith and Brady, 1997) with spatial extent threshold of 4 mm, and temporally high pass filtered (pass band > 0.01 Hz) prior to single-subject ICA for each dataset. Spatial ICA was performed using FSL MELODIC. The analysis was constrained to obtain only 80 components for each subject to limit the processing time. This number was found to be liberal enough to reliably identify the white matter component. The white matter component was identified in each subject by an experienced observer (SK) in all but three subjects. Data for these three subjects was observed to be high in temporal noise and multi-band artefacts (between slice signal correlations).

### 7 Tesla MRI venography and BOLD imaging

To confirm the venous origin of the white matter IC map, a single healthy volunteer (26 year old female with no history of neurological or vascular disease) was imaged using a 7 Tesla MRI System (Magentom, Siemens, Erlangen). Two sets of images were collected: (1) a gradient echo sequence for calculation of the susceptibility weighted image (repetition/echo times = 24/17.34 ms; flip angle = 13°; resolution 0.75×0.75×0.75 mm^3^ axial acquisition); and (2) a multi-band BOLD-weighted echo planar imaging sequence with imaging parameters comparable to those used for 3T MRI (repetition/echo times = 1500/25 ms; flip angle = 44° resolution = 2×2×2 mm^3^ axial acquisition; number of BOLD measurements = 205), acquired while subjects watched a blank screen with eyes open.

BOLD data were processed using MELODIC according to the methods described above for 3 Tesla data and linearly registered to the high-resolution gradient echo image using FLIRT (FSL, FMRIB, Oxford, UK).

### White matter venous hypercapnia challenge

To test the effects of non-neurally driven BOLD signal changes on the white matter venous system, three healthy male subjects underwent two runs of functional MRI whilst performing a simple breath hold task to induce transient hypercapnia. The first run consisted of 7 repetitions of 18 sec breath hold followed by 24 sec normal breathing. The second run consisted of 24 sec of breath hold followed by 18 sec of normal breathing. BOLD functional MRI data were preprocessed to correct for motion, high-pass and spatially filtered as previously described above. Mixed-effects general linear models were used to test for the following main effects: condition (hypercapnia vs baseline), tissue type (grey vs white matter) and duration (short vs long breath hold), and following interactions: condition/tissue type and condition/tissue type/duration. Post-hoc correlation analyses were used to compare BOLD signal changes between tissue types.

### White matter venous BOLD spectral analysis

In order to compare the power spectra of actual BOLD signal time-courses between patients and controls, rather than IC time-courses, the white matter probability map was used to calculate the weighted-average power-spectrum for all brain voxels for each subject. Briefly, raw BOLD time-course data were pre-processed to correct for head motion and perform high pass filtering. No spatial smoothing was performed. For each voxel, the BOLD time-course was converted to a power spectrum using nonparametric multi-taper spectral decomposition (Thomson, 1982) implemented in the MATLAB^®^ R2016a Signal Processing Toolbox (mathworks.com/help/signal/ref/pmtm.html). A weighted mean power spectrum was calculated for each subject using the weighting values from the spatial probability map. The average power within a conservatively judged peak region (0.04 to 0.1 Hz) was compared between patients and control subjects using a Student’s t-test. Pearson’s correlation analyses were performed between average power and logarithmically transformed lesion volume and brain, grey and white matter fractions.

### White matter venous BOLD voxel wise group comparisons

A data driven signal modelling approach was used to test for putative anatomical differences in the white matter BOLD signal between patients and controls. For each brain voxel, high-pass filtered BOLD signals were converted to power spectra using multi-taper spectral decomposition in MATLAB® R2016a. Each voxel’s power spectrum was fitted with a Gaussian curve using nonlinear least squares with appropriate boundary conditions (0.04<*μ*<0.1 Hz; *σ*>0 Hz) using MATLAB® R2016a. The fitted power and frequency for the spectral peak was compared voxelwise between groups using a general linear model permutation sampling method (RANDOMISE, FSL, FMRIB, Oxford, UK) (Winkler et al., 2014) family-wise error corrected with threshold-free cluster enhancement (Smith and Nichols, 2009).

## Results

### Characterisation of the white matter BOLD signal in healthy subjects

Spatial ICA is a common method for identifying independent neural networks in resting-state fMRI. Previous observations of a component within white matter that appeared to co-localise well with the known anatomy of the internal cerebral venous system (Salimi-Khorshidi et al., 2014) led us to examine this signal specifically in a group of 14 healthy individuals using 3 Tesla MRI. Functional MRI data were collected at rest and pre-processed (corrected for head motion, high pass filtered and spatially smoothed) before analysis with ICA (Beckmann and Smith, 2004). In all subjects a component was identifiable in the periventricular white matter (Fig. 1A). Single sample statistical analysis demonstrated a large region after family wise error correction across most of the cerebral white matter (Fig. 1B).

**Fig. 1.**
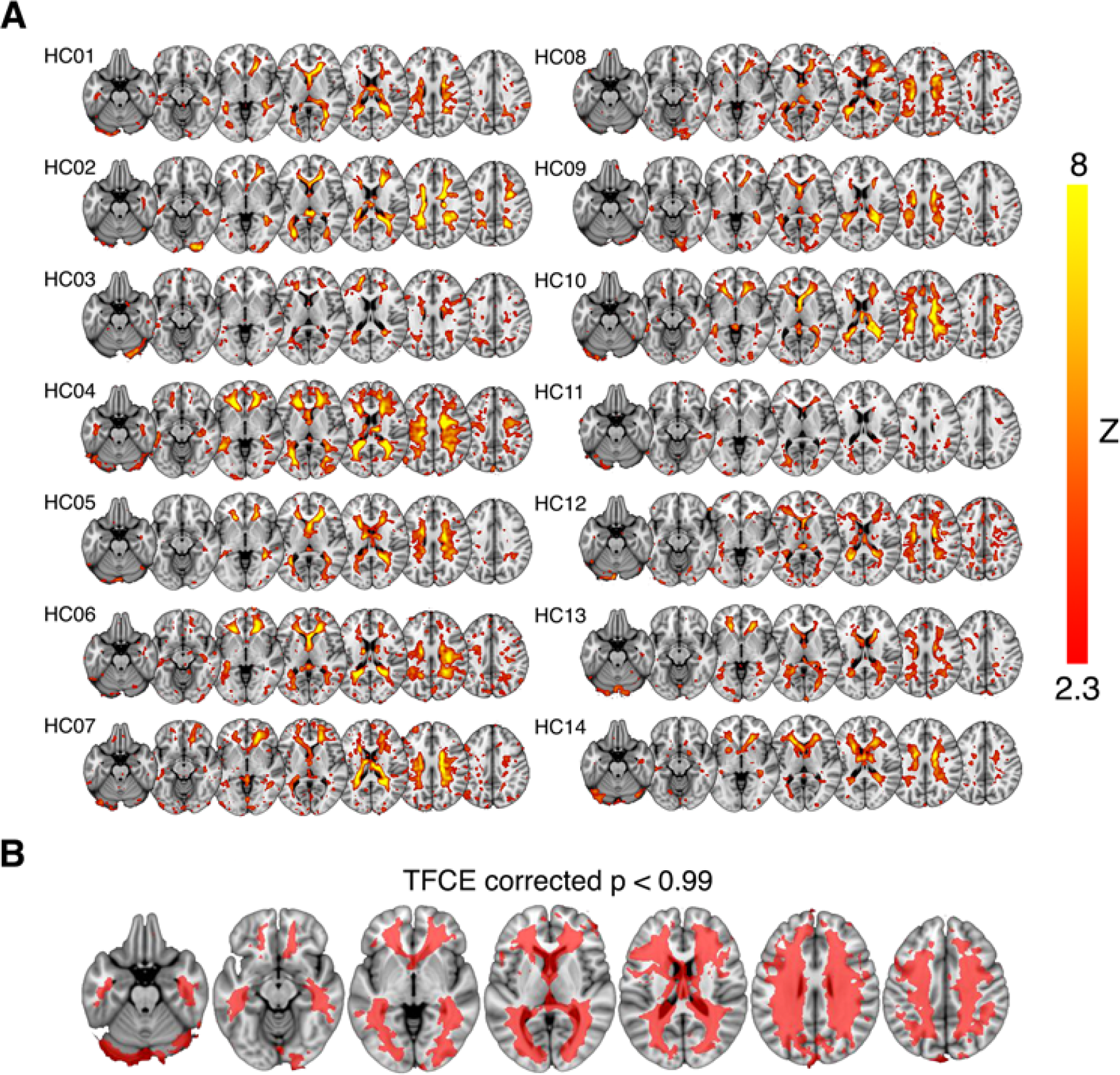
Spatial maps of the white matter BOLD signal output from spatial independent components analysis. (A) Shows individual component maps for the white matter BOLD signal. (B) Shows the regions of significant correspondence across subjects using a single group t-test, family-wise error corrected using the threshold-free cluster enhancement (TFCE) method.

The anatomy of the signal was investigated more precisely in a single subject using high-resolution functional and susceptibility-weighted MRI at 7 Tesla (Fig. S1). Susceptibility weighted imaging reveals the venous blood vessels within the white matter as regions of high magnetic susceptibility due to the presence of blood. The ICA component was most evident in regions characterised by medium to large periventricular veins.

Power spectral features of the white matter BOLD time-series were also assessed. In all subjects, white matter BOLD signals were characterised by a narrow peak frequency band was observed in the range of 0.05 to 0.070 Hz [mean (SD) = 0.061 (0.008) Hz; peak power (SD) = 34.87 (7.20) dB/Hz]. Peak power and frequency were not correlated (R=−0.28).

### Replication analysis of the venous BOLD signal using HCP dataset

To confirm the spatial and spectral features of the white matter BOLD signal, we replicated the previous analyses in an independent dataset of 30 healthy resting state fMRI datasets obtained from the Human Connectome Project (HCP) (Van Essen et al., 2013). Compared to the mean white matter IC map for our dataset, the HCP dataset displayed lower a lower average magnitude (Fig. 2A). The same narrow peak frequency band was observed in the HCP data and did not differ significantly in peak frequency or power from our dataset [mean frequency (SD) = 0.057 (0.008) Hz, *t*_33_ = 1.36, *p* = 0.20; mean peak power (SD) = 32.09 (6.12) dB/Hz, *t*_33_ = 1.33, *p* = 0.21] (Fig. 2B). There was no correlation between peak power and frequency (R = −0.23).

**Fig. 2.**
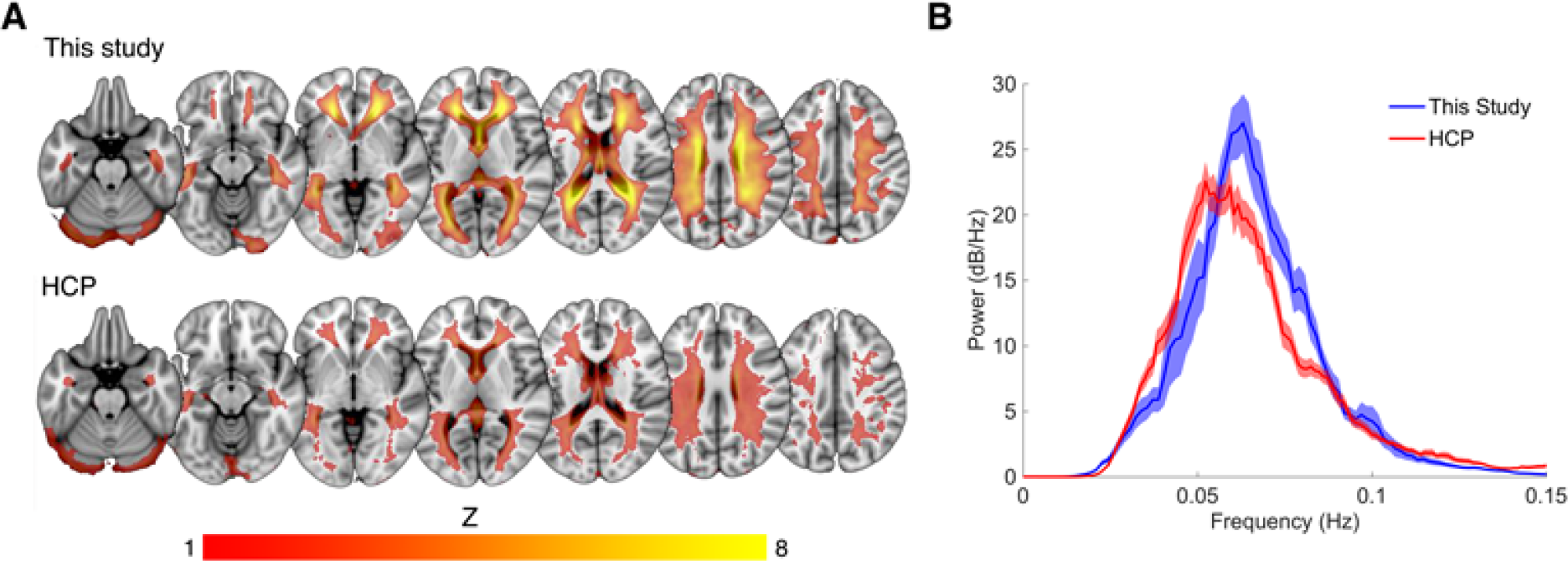
Comparison of spatial and spectral characteristics for the white matter BOLD signal between this study the Human Connectome Project (HCP). (A) Shows the high degree of spatial congruence between the mean component maps of the two data sets. (B) The spectral properties of the venous BOLD signal were similar between the datasets, with both showing a narrow peak high power band between 0.04 and 0.1 Hz.

The white matter component was undetectable in three HCP subjects. To explore this further we compared temporal SNR between our data and the HCP data to determine whether temporal noise could reduce the detectability of the venous component. We observed a relatively large decrease in tSNR in the HCP data compared to our data ~60%, despite the large number of extra fMRI volumes (n = 400 for our data compared to n = 1200 for HCP data). This difference was likely due to a combination of smaller voxel dimensions (our data = 2.5mm isotropic, HCP = 2mm isotropic) and higher multiband acceleration factor (our data = 3, HCP = 5) used by HCP. However, despite noisier and thus less reliable data, the HCP data recapitulated the gross spatial and temporal features of the venous BOLD signal observed in our dataset.

### Venous BOLD physiology during hypercapnia

To better understand the physiological mechanisms driving the white matter BOLD oscillation, we employed dynamic hypercapnic challenge (breath holding) during fMRI scanning in three healthy subjects. Hypercapnia is a potent vasodilator in cerebral grey matter, causing marked BOLD signal attenuation in the cortex commensurate with increased partial volume inclusion of low contrast blood. We therefore expected that hypercapnia induced vasodilation would also affect the venous BOLD signal. Mixed-effects general linear models were used to test for the following main effects: condition (hypercapnia vs baseline), tissue type (grey vs white matter) and duration (short vs long breath hold), and following interactions: condition/tissue type and condition/tissue type/duration. Only two contrasts were significant. Firstly, we observed a significant main effect of condition (F_1948,1_=53.8, p<0.0001) demonstrating a clear effect of hypercapnia on the BOLD signal. Secondly, we detected a significant interaction between condition and tissue type (F_1948,1_=1269, p<0.0001) demonstrating that grey matter and white matter responded differently to hypercapnia. Post-hoc comparisons of BOLD change between grey and white matter showed that while grey matter BOLD reduced as expected (Δ marginal means =−1.50, p<0.0001), white matter BOLD signals significantly increased (Δ marginal means = 0.98, p<0.0001). The degree of BOLD increase in the white matter was negatively correlated with grey matter BOLD decreases in all three subjects for both short (subject 1: R=−0.44, p<0.0001; subject 2: R=−0.73, *p*<0.0001; subject 3: R=−0.50, *p*<0.0001) and long (subject 1: R=−0.35, *p*<0.0001; subject 2: R=−0.63, *p*<0.0001; subject 3: R=−0.46, *p*<0.0001) breath hold runs (Fig. S2). Together, these data suggest that the white matter BOLD signal is associated with expansion and contraction of the vessel walls that can be modulated by hypercapnia. We hypothesise that the unforeseen increase in white matter BOLD signal (vasoconstriction) during hypercapnia acts to maintain intracranial pressure associated with cortical and subcortical vasodilation. This is consistent with the well-recognised role of veins as blood capacitance vessels (Auer and Loew).

### White matter BOLD alterations in early multiple sclerosis

A relatively homogeneous cohort of 18 multiple sclerosis cases was selected for comparison. All patients were diagnosed with multiple sclerosis diagnosed based on the 2010 McDonald criteria (Polman et al., 2011) and all had presented with acute optic neuritis 3-5 years previously. One patient was excluded due to excessive head motion during the resting fMRI scan.

To compare patients and controls, we first created an unbiased template of the white matter venous system based on ICA outputs from all healthy and multiple sclerosis subjects in our study. Individual subject white matter IC maps were thresholded to remove voxels with low Z scores (Z<2.3), binarised and transformed to standard MNI-152 brain space using nonlinear spatial registration. This yielded a standard space white matter venous map with voxel values ranging from 0 (white matter component present in no subjects) to 1 (white matter component present in all subjects) (Fig. 3A). The template was used to calculate the weighted average raw BOLD signal time-courses for the white matter for each subject (voxels within T2 lesions in patients were omitted in patients). These time-courses were high pass filtered (pass band > 0.03 Hz) to remove low frequency artefacts (e.g. signal drift due to MRI gradient heating), and spectrally analysed using the multi-taper spectral decomposition (Fig. 3B and C).

Average white matter BOLD power across a conservatively judged peak frequency range (0.04 to 0.1 Hz) was significantly lower in multiple sclerosis patients (6.70±0.94 dB/Hz) compared to control subjects (7.64±0.71 dB/Hz, t_28_ = 2.9, *p* = 0.002) when compared using a *t*-test (Fig. 3D). Variation in the mean power in this band in patients correlated significantly with logarithmically transformed lesion volume (R=−0.65, *p*=0.005) (Fig. 3E), but not with non-inflammatory markers of brain injury including normalised brain (R = 0.22), grey matter (R = 0.31) or white matter volumes (R = 0.33). These overall statistical results were not affected by narrowing the frequency band used for calculating mean power.

**Fig. 3.**
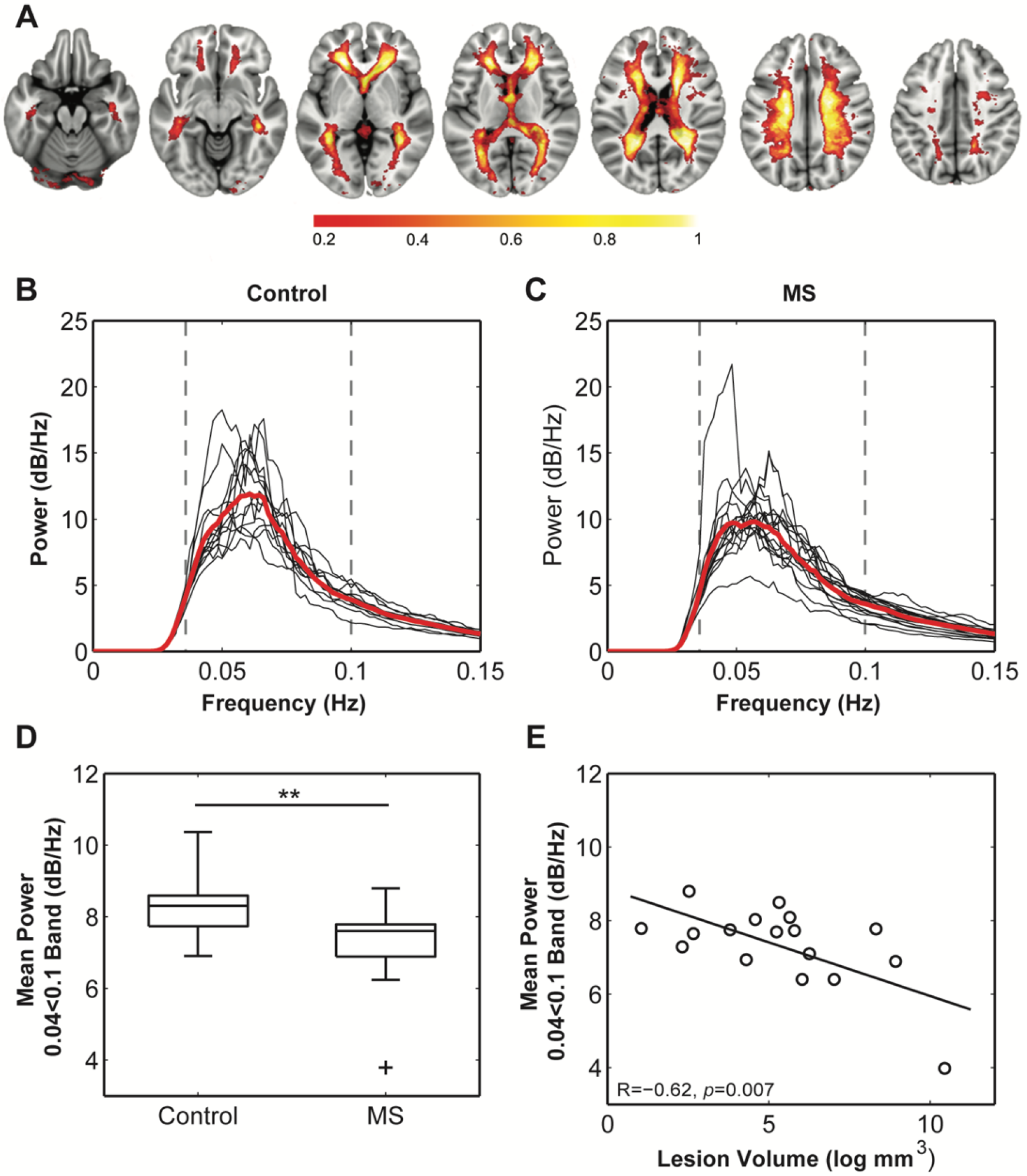
Comparison of the white matter BOLD signal between healthy and multiple sclerosis participants. (A) The probabilistic map of the BOLD signal across all subjects. Values (0 to 1) reflect the proportion of subjects with the component in each voxel. Maximal congruence was observed in regions corresponding to the location of the peri-ventricular veins and small veins within the corona radiata. These are also the most common sites of neuro-inflammatory activity in multiple sclerosis. (B) and (C) Weighted average power spectra from within the probability map for control and multiple sclerosis subjects demonstrate a clear peak power within the 0.04 to 0.1 Hz frequency band. (D) Mean power within this band was significantly reduced in multiple sclerosis cases compared to control subjects (** = p<0.01), and (E) mean power in patients significantly correlated with logarithmically transformed cerebral lesion volume.

### BOLD power spectrum modelling

Finally, we used a spatially unbiased method to compare white matter BOLD signals between healthy and multiple sclerosis subjects based only on the identified spectral characteristics (narrow peak power in 0.04 to 0.1 Hz frequency range). Raw BOLD time-courses for every brain voxel in each subject were high pass filtered (<0.03 Hz) and converted to frequency domain using multi-taper spectral decomposition. For each voxel, a Gaussian curve was fitted to the frequency spectrum using nonlinear least squares with appropriate boundary conditions (0.04<*μ*<0.1 Hz; *σ*>0 Hz). This yielded voxel-wise estimates of peak power in the relevant frequency range. In healthy subjects, peak power was constricted to the white matter (Fig. 4A top) demonstrating the specificity of the spectral characteristics to the white matter. Similarly, the multiple sclerosis group showed peak power constrained to the white matter, yet the magnitude of the power was diminished compared to control (Fig. 4A bottom). Voxel-wise statistical analyses revealed significant loss of power in the periventricular white matter (Fig. 4B), consistent with the location of multitudinous small veins that are a common site of inflammatory demyelination in multiple sclerosis.

**Fig. 4.**
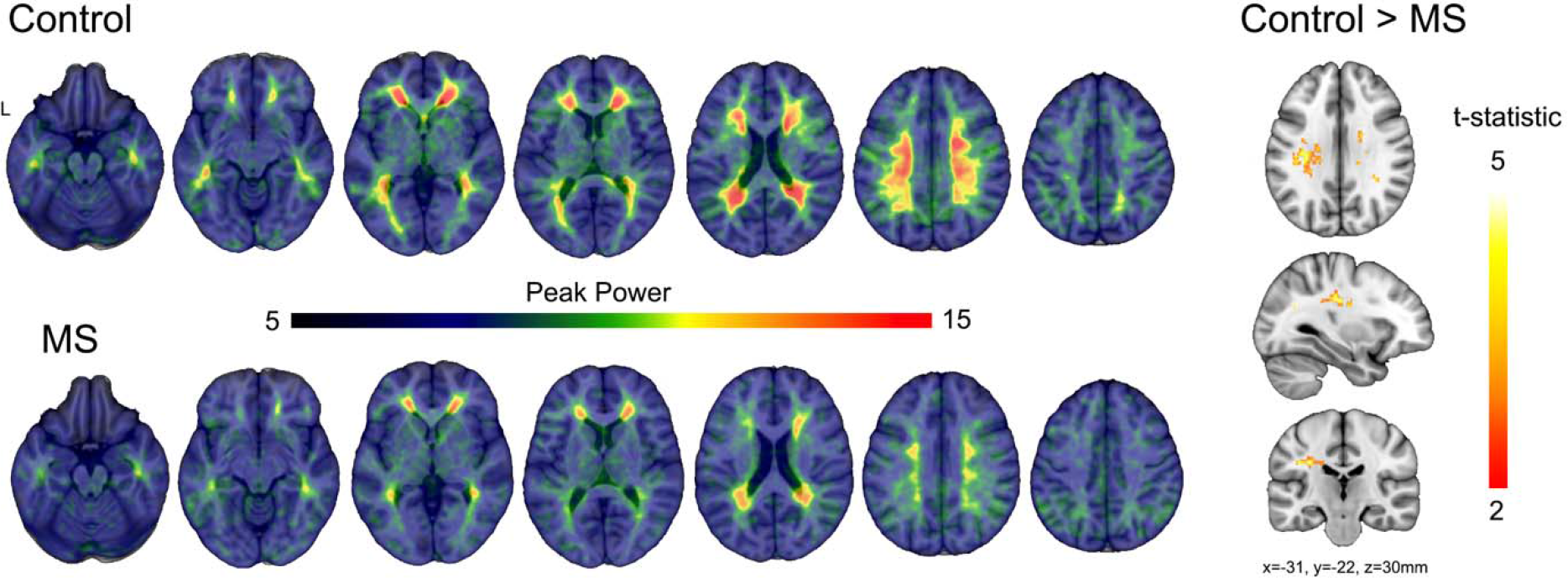
Comparisons of modelled BOLD power between multiple sclerosis and control subjects. (A) Average modelled peak power of the white matter BOLD signal for control and multiple sclerosis subjects illustrates a reduction in peak power (dB/Hz) across the much of the white matter. (B) The t-statistic map shows regions of significant reduction in peak power in the multiple sclerosis group compared to control subjects after correction using threshold-free cluster enhancement.

## Discussion

This study represents, to the best of our knowledge, the first in-depth study of the haemodynamics of the white matter using BOLD-weighted MRI. The white matter BOLD signal has been reported previously in the context of removal of artefactual signals from resting state neural connectivity analysis [see Fig. 3 from Salimi-Khorshidi et al., (2014)]. Perivenular lesions are a consistent feature of multiple sclerosis (Tallantyre et al., 2011), and the high degree of spatial congruence between the white matter BOLD signal and regions of high lesion load in multiple sclerosis (Vellinga et al., 2009) led us to explore the signal further as a potential marker for venous pathophysiology. We confirmed that the white matter BOLD signal co-localised with smaller and larger internal cerebral venous system using high resolution susceptibility-weighted imaging at 7 Tesla. In our experience, white matter BOLD signals are almost always observed in single subject ICA output but are generally not analysed. Our analyses confirmed that the white matter BOLD signal is highly stereotypic across individuals in terms of anatomy and spectral properties. We were also able to confirm spatial and temporal features of white matter BOLD in a completely independent dataset (HCP). We expect that the signal could be obtained from any resting-state fMRI data given the reasonably low frequency of the signal (~0.05 Hz) compared to the acquisition frequencies used in most fMRI studies (0.33 to 1.3 Hz).

While we are not able to provide a definitive physiological account for the white matter signal using *in-vivo* imaging data, we were able to physiologically manipulate the signal using a transient hypercapnia challenge. Hypercapnia caused a transient increase in the white matter BOLD signal. A likely cause for this increase is a reduction in the voxel partial volume inclusion of low contrast venules and veins associated with vasoconstriction. Vasoconstriction in the white matter venous system under hypercapnia could be a mechanism for maintaining intracranial pressure during the large vasodilatory response in grey matter regions characteristic of hypercapnia.

At rest, the white matter BOLD signal was observed to oscillate at around one cycle every 20 seconds. This rate excludes cardiac and respiratory influences and instead points to a potential auto-regulatory function. Cerebral blood flow is tightly regulated and is resistant to rapid changes in systemic arterial blood pressure (Lassen, 1964). The observed BOLD frequency range is within the limits of both myogenic (0.02 to 0.15 Hz) and sympathetic (0.07 to 0.15 Hz) vascular regulation (Stauss, 2007). In support of a myogenic mechanism, mechanoreceptors have been identified in the walls of the cerebral venous system that are activated by an increase in cerebral blood volume to regulate blood flow (McHedlishvili, 1980). The degree of correlation of myogenic regulation across the venous system is unknown. In contrast, sympathetic activity is centrally controlled and known to control vasoconstriction within the cerebral venous system (Auer et al., 1981). Given the high degree of signal coordination across the entire internal cerebral venous system in healthy individuals, we hypothesise that the observed oscillatory BOLD signal is likely to reflect cyclic vasodilation and constriction under central sympathetic control. Further studies will be required confirm the sympathetic origin of the signal and to elucidate the local control mechanisms.

In multiple sclerosis patients, power in the white matter BOLD signal was reduced commensurate with the degree of neuroinflammatory lesion load. This suggests that the physiological substrates of BOLD signal power loss in people with multiple sclerosis relates to inflammatory damage. The white matter venules and veins are the principal entry point for peripheral immune cells into the brain through a disrupted blood-brain barrier during acute inflammation. Such damage to the vascular wall acutely could lead to ongoing postacute damage or imperfect repair. Indeed, previous studies have reported venous atrophy (Ge et al., 2009; Sinnecker et al., 2013; Zivadinov et al., 2011) and altered blood flow (Varga et al., 2009) in the white matter of people with multiple sclerosis. Three studies have also reported reduced venous density in multiple sclerosis (Ge et al., 2009; Sinnecker et al., 2013; Zivadinov et al., 2011), most likely associated with venous atrophy to the point where the blood vessels can no longer be visualised, even with the high resolution (<=0.5×0.5 mm^2^ in-plane) afforded by 7 Tesla MRI (Sinnecker et al., 2013). Consistent with our findings of a link between neuroinflammation and venous damage, Sinnecker and colleagues (2013) showed that venous density was negatively correlated with T2 lesion load. Venous atrophy is a potential substrate for loss of BOLD signal power via the reduction of partial volume inclusion of the venous signal in imaging voxels. Also potentially associated with venous atrophy, reduced venous flow has been reported in the periventricular white matter in both clinically isolated syndromes and relapsing-remitting multiple sclerosis patients using dynamic contrast MRI (Varga et al., 2009). The authors also observed a non-significant trend towards reduced blood volume, suggestive of vascular atrophy. Gaitán and co-workers (2013) reported narrowing of veins within T2 lesions, yet enlargement of perilesional veins compared to healthy control veins using ultra-high resolution T2*-weighted MRI (0.5×0.5×0.5 mm^3^) during gadolinium infusion. The authors interpreted their findings to indicate that perivascular inflammation could lead to vascular compression and thickening of the perivascular wall. Supporting evidence can be found in histological studies of the vascular system in multiple sclerosis noting fibrinoid and haemosiderin deposition, thrombosis and venous wall thickening (Adams, 1988; Adams et al., 1985). Given the large capacitance of the venous system, it is possible that the perilesional enlargement observed by Gaitán and coworkers (2013) reflects a bottleneck effect caused by flow resistance within lesions, leading to perilesional vascular distension. Together, these studies demonstrate significant neuroinflammatory damage to the white matter venous system that could account for the haemodynamic changes observed in our study. It is conceivable that reduced haemodynamics has the potential to exacerbate neural damage via local ischemia. This hypothesis support further investigation of the venous BOLD signal in the context of other diseases characterised by white matter lesions.

This study has several limitations that should be addressed in follow-up studies. Firstly, we did not directly anatomically image and map the venous system in all subjects, so it was not possible to compare the haemodynamic alterations to venous anatomy and morphology directly. The high resolution afforded by high field (7T+) MRI allows the mapping of fMRI signals to specific blood vessels, and to create maps of the white matter venous system that could be used to better characterise the spatial distribution of venous damage. We did however, exclude voxels from each patient’s white matter venous probability map that were classified as lesion on double inversion recovery scans. Therefore, the changes in white matter BOLD power observed in patients was measured from normal appearing brain regions that were not influenced by overt signal changes associated with lesion pathology. Finally, our study focussed on a convenience sample of patients with a history of acute optic neuritis with a relatively consistent and short disease duration of between 3 and 5 years. Future studies should characterise the progression of white matter haemodynamic abnormalities in later disease stages. Longitudinally designed studies will also be required to determine whether white matter haemodynamic abnormalities are associated with greater susceptibility to subsequent inflammatory cell infiltration or neurodegenerative changes.

### Summary and Conclusions

The internal cerebral veins within the white matter are the most common site of early peripheral immune cell infiltration in the neuroinflammatory demyelinating disease multiple sclerosis. This study identified a novel haemodynamic signal with a narrow spectral peak in the cerebral veins of the white matter using resting-state functional MRI and ICA. Peak power of the signal was reduced in people with multiple sclerosis, and the degree of reduction correlated significantly with neuroinflammatory lesion volume. These results indicate that multiple sclerosis is associated with dysfunction of the white matter veins that initiates early in the disease. Future studies are required to identify the physiological mechanism driving white matter BOLD haemodynamics and explore the role of altered haemodynamics in multiple sclerosis pathophysiology.

## Acknowledgments

We sincerely thank all study participants for their time.

## Author contributions

SK conceived the study, collected and analysed data and wrote the manuscript, SG analysed data and edited the manuscript, JC analysed data and edited the manuscript, TK conceived the study and edited the manuscript

## Data and materials availability

Raw anonymised data can will be made available upon request.

---

**Fig. S1.**
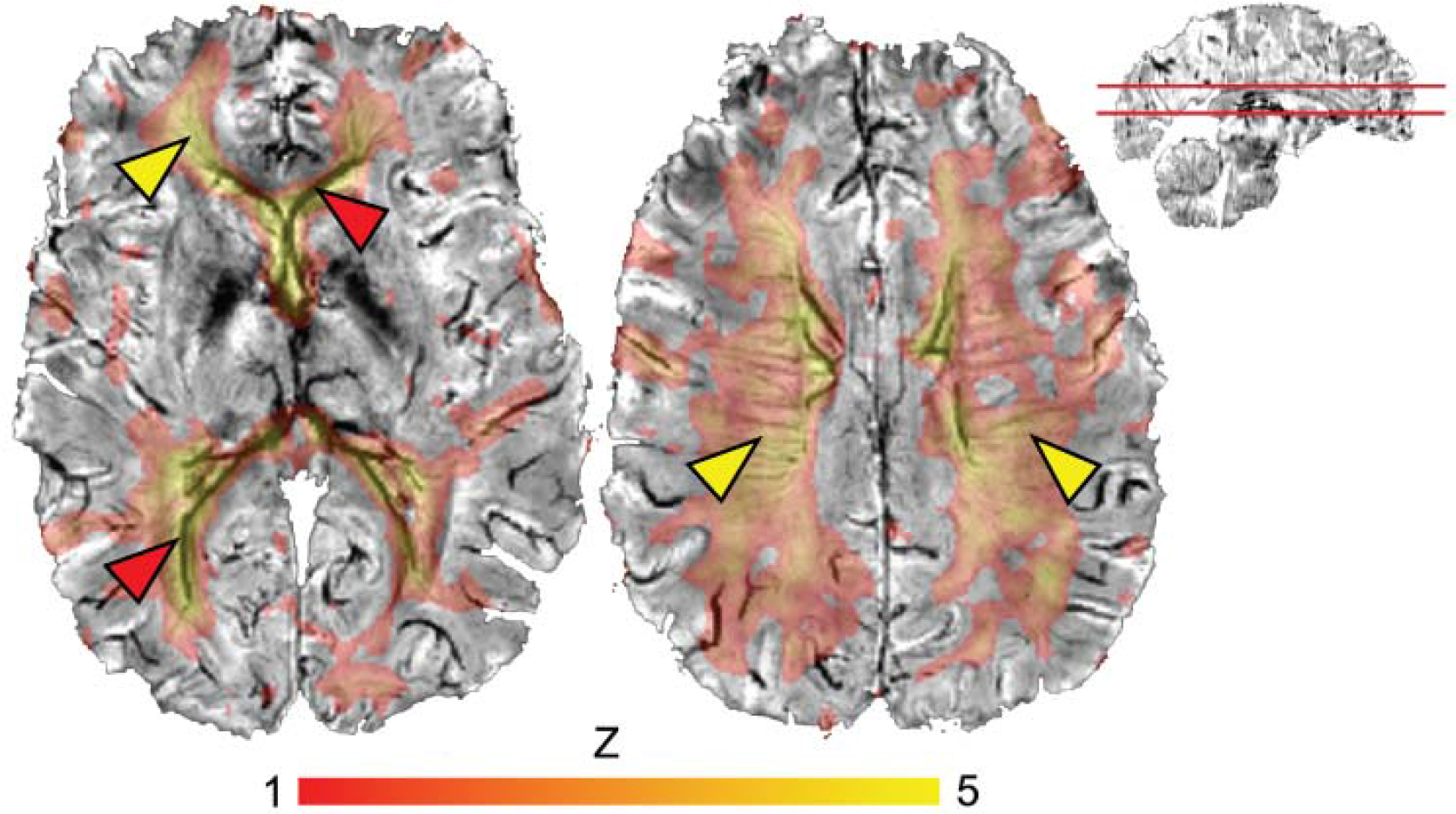
Venous BOLD signal ICA output overlayed on the same subject’s susceptibility weighted MRI scan. Veins appear as dark lines in the susceptibility-weighted image, and these co-localise with regions of highest venous BOLD signal in both smaller veins deep in the white matter (yellow arrowheads) and larger periventricular veins (red arrowheads).

**Fig. S2.**
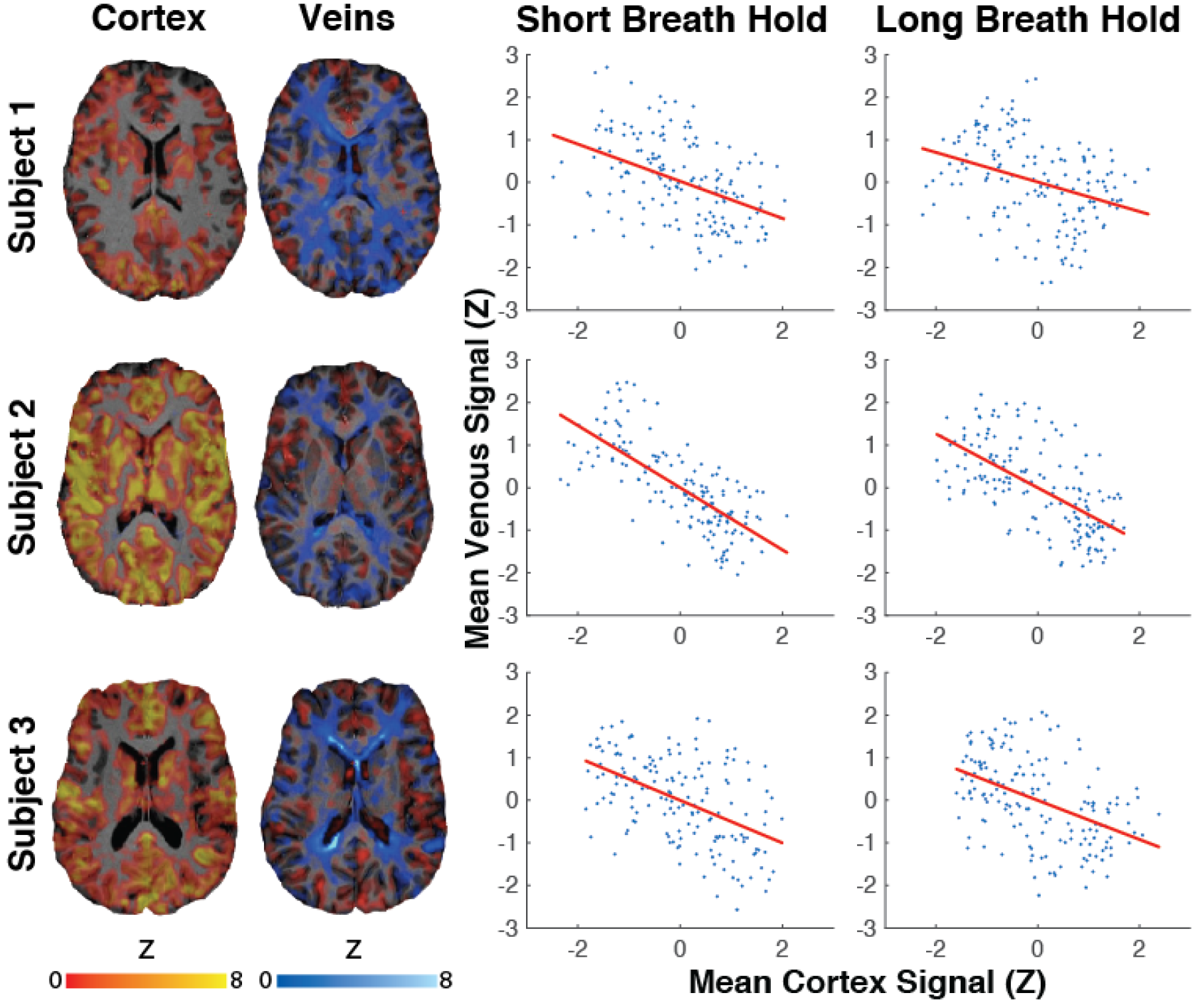
Comparison between BOLD signals from cortex and internal cerebral veins during transient hypercapnia. (breath hold) in three healthy subjects. The left side shows the spatial maps for cortex and white matter that displayed negative and positive signal changes respectively during breath holds. Signal time course correlations across the entire experiment (right) show that the decrease in cortical BOLD was associated with a contemporaneous increase in venous BOLD.

